# Maternal antibodies to gliadin and autism spectrum disorders in offspring - A population-based case-control study in Sweden

**DOI:** 10.1101/2020.02.13.948620

**Authors:** Renee Gardner, Ida Samuelsson, Emily G. Severance, Hugo Sjöqvist, Robert H. Yolken, Christina Dalman, Håkan Karlsson

## Abstract

**Objective:** Individuals diagnosed with autism spectrum disorders (ASD) are reported to have higher levels of antibodies directed towards gliadin, a component of wheat gluten. However, no study has examined such antibodies in etiologically-relevant periods before diagnosis. The objective of this study is to investigate if maternal levels of immunoglobulin G antibodies directed at gliadin, during pregnancy and at the time of birth, are associated with ASD in offspring.

**Methods:** In this population-based study set in Sweden with 921 ASD cases and 1090 controls, we analyzed levels of anti-gliadin antibodies (AGA) in archived neonatal dried blood spots (NDBS, as maternal IgG is transferred to the fetus) and in paired maternal sera collected earlier in pregnancy for a subset of 547 cases and 428 controls. We examined associations to any ASD diagnosis and considering common comorbidities (i.e. intellectual disability [ID] and attention-deficit/hyperactivity disorder [ADHD]). We compared 206 ASD cases to their unaffected siblings to examine the potential for confounding by shared familial factors.

**Results:** High levels (≥90^th^ percentile) of maternal AGA were associated with decreased odds of ASD, particularly ASD with comorbid ID, when measured in NDBS (OR 0.51, 95% CI 0.30–0.87) with a similar trend in maternal sera (0.55, 0.24-1.29). High levels of maternal AGA were similarly associated with lower odds of ASD with ID in the sibling comparison.

**Conclusions:** This first study of exposure to AGA in the pre- and perinatal periods suggests that high levels of maternal AGA are associated with lower odds of ASD with ID.

## Introduction

Individuals with Autism spectrum disorders (ASD) are diverse in their presentation of core symptoms (1), and are at increased risk for both psychiatric and somatic comorbidities (2–4). For instance, about one third also present with intellectual disability (ID), and ADHD respectively (3, 5). The clinical characterization of ASD as a combined diagnostic group, might thus not reflect one common, but possibly several distinct, underlying pathogenic processes that remain a challenge for research that aims to identify the etiology of ASD.

Moreover, a genetic overlap between psychiatric and immunological disorders including autoimmune diseases have been established (4, 6). Celiac disease (CD), an autoimmune disorder triggered by gluten exposure, has been discussed in connection to ASD for over 40 years based on the observation that gastro-intestinal symptoms are common in individuals with ASD (7). However, studies have been inconsistent. While some studies report increased CD prevalence among individuals with ASD (8), other studies find no such associations (9–11). Several studies report children with ASD to have significantly higher levels of antigliadin IgG antibodies (AGA) compared to controls (12–14) in the absence of more specific serological markers of CD, such as IgA anti-transglutaminase-2 and anti-endomysial antibodies (12). Whether such reactivity against gluten is causally involved in ASD is, however, not known as previous studies have focused on individuals already diagnosed with ASD. Recent dietary intervention studies indicate a lack of efficacy of gluten-free diets in the treatment of ASD (15–17).

The objective of the present study was to investigate if maternal levels of IgG antibodies directed at gliadin, measured during pregnancy and in the neonate during the first days of life, are associated with a later diagnosis of ASD. IgG detected in the neonatal circulation is almost exclusively transferred from the mother through the placenta, and we consider AGA measured in neonatal and maternal samples to be of maternal origin. Comparisons within sibling-pairs discordant for ASD were conducted to address the potential for residual confounding by shared familial factors.

## Materials and Methods

### Study population

This study is nested within the Stockholm Youth Cohort (SYC), a register-based cohort of all children aged 0-17 years living in Stockholm County (18, 19). We selected all individuals born 1996-2000 and diagnosed with ASD as of December 31, 2011, a random sample of the 98,597 individuals in the SYC born 1996-2000, and unaffected siblings to ASD cases born 1996-2000 (Figure 1). The 1996-2000 birth cohorts were chosen to balance the (time-dependent) quality and availability of archival biological specimens with sufficient follow-up time in registers to maximize chance of a registered diagnosis of ASD. Diagnostic information in the SYC was updated after sample collection to include follow-up until December 31, 2016.

**Figure 1.**
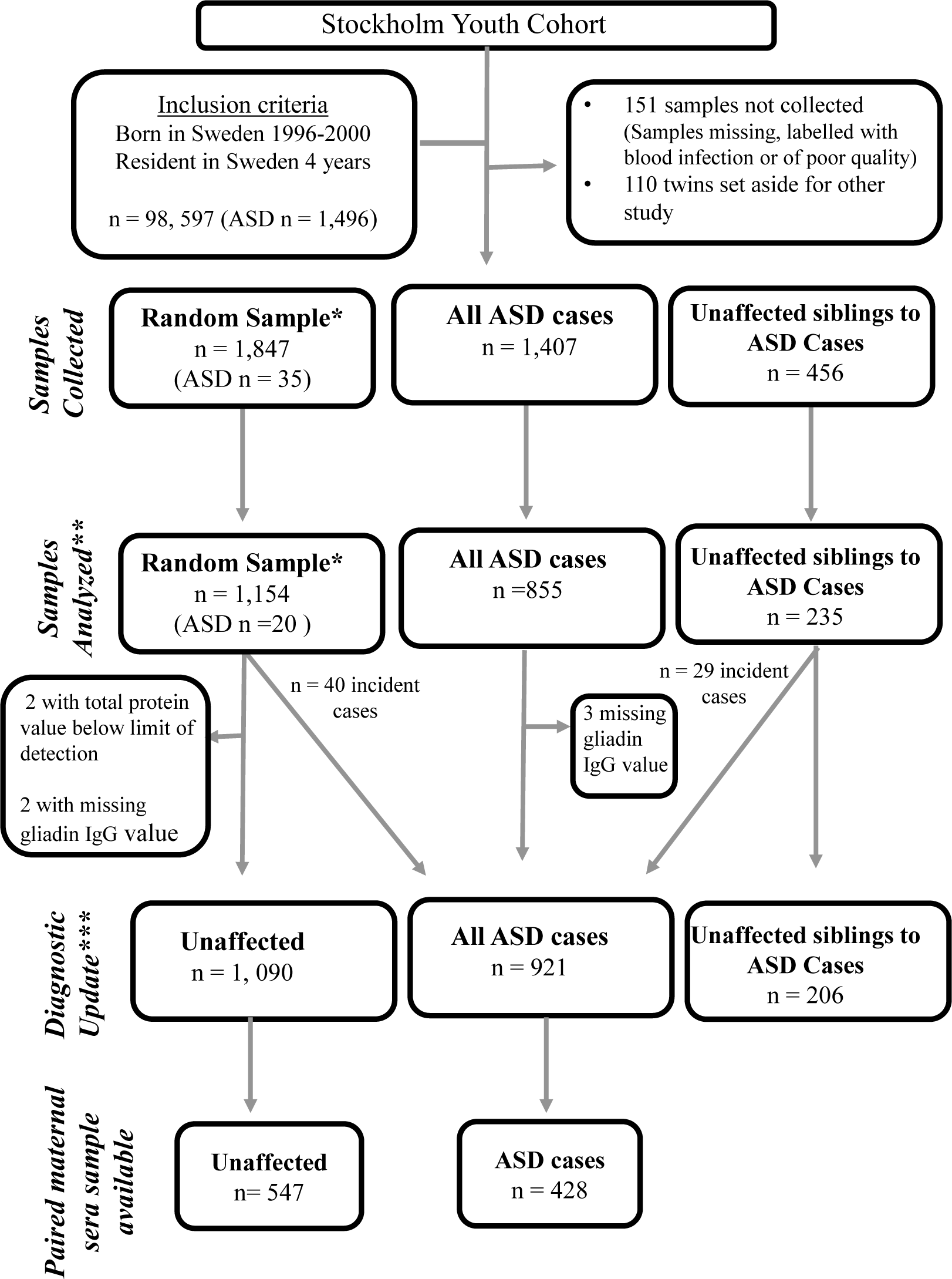
Flow chart describing sample selection and study populations. *A random sample of the SYC and, as such, this includes ASD cases also included in the total sample of all ASD cases ** Smaller samples to be analyzed were sampled out from each group due to economic reasons, based on birth year. To maximize chance of available paired maternal sera the birth years 98 00 were favored, and approximately 12 of individuals born 96 97 were included. ***ASD diagnostic data updated as of December 31, 2016 for the SYC, resulting in 69 incidence cases of ASD: 29 individuals sampled as siblings to an ASD case and 40 individuals sampled as part of the random sample of the SYC. Abbreviations: ASD = Autism spectrum disorders, NDBS = Neonatal dried blood spots

Ethical approval was provided by the Stockholm regional review board (DNR 2010/1185-31/5). Individual consent was not required for this anonymized register-based study.

### ASD Ascertainment

ASD case status as of December 31, 2016 was ascertained according to a validated case-finding approach, described in detail elsewhere (19). See ST1 for all diagnostic codes used for case ascertainment. In line with recommendations from DSM-V (1), we studied ASD as a combined diagnostic group. ASD outcomes were also stratified by absence or presence of comorbid ID and ADHD into three mutually-exclusive diagnostic categories; ASD with ID, ASD with ADHD and ASD “only” (indicating an ASD diagnosis without comorbid ID or ADHD). Individuals with both comorbid ID and ADHD were included in the ASD with ID group, as we considered ID to be the more severe disorder.

### Biological Samples: Neonatal dried blood spots

Archival NDBS were collected from the national biobank, Centre For Inherited Metabolic Diseases, Karolinska University Hospital, Solna. The mean age (±SD) of the neonate at blood sampling was 4.07 ± 1.25 days. A 3.2 mm diameter disc was punched from each blood spot and distributed into deep 96-well plates. Proteins were eluted from the filter paper by incubation in 200 microliters of phosphate-buffered saline for 2 hours on a rotary shaker (600 rpm) at ambient temperature. Total protein content in each eluate was measured using infra-red spectroscopy (Direct Detect, Merck Millipore, Burlington, MA, USA). The remainder was aliquoted and stored at −80 °C.

### Biological Samples: Maternal sera

Maternal sera from the same pregnancy, originally collected during pregnancy as part of regional screening programs at a mean gestational age (±SD) of 11.97 ± 5.02 weeks, were retrieved from regional biobanks in Stockholm County for a subset of the study individuals (see Figure 1). Maternal sera samples were thawed and an aliquot was transferred to a new tube and sent to us on dry ice. These samples were thawed, re-aliquoted and immediately frozen at −80 °C.

To maximize chance of an available maternal sera from the same pregnancy, we primarily selected individuals in the 1998-2000 birth cohorts for the laboratory analysis of neonatal dried blood spot (NDBS) samples (Figure 1). In order to analyze time-dependent influences, a small sample (12.5%) of those born 1996-1997 was also selected. Our final sample with analyzed NDBS consisted of 2011 individuals (1090 controls and 921 ASD cases) and 206 unaffected siblings. For 975 of these individuals, archival paired maternal sera obtained earlier in that same pregnancy were available, with a final sample with matched maternal sera and NDBS analysis of 547 controls and 428 ASD cases.

### Laboratory Analysis of AGA

NDBS eluates (diluted 1:20) and maternal sera (diluted 1:100) were analyzed using commercially available anti-gliadin IgG enzyme-linked immunosorbent assays (ELISA) according to manufacturer’s protocols, (IBL America, Minneapolis, Minnesota, U.S.A.). All plates were read in a microtiter plate reader at 450 nm and optical density values were used in statistical analyses.

### Maternal levels of total IgG

We analyzed maternal sera (diluted 1:250,000) for total levels of IgG antibodies. Commercially available ELISAs for quantitative detection of human total IgG were used according to manufacturer’s protocols (Thermo Fisher Scientific, Waltham, MA, U.S.A.). All plates were read in a microtiter plate reader at 450 nm and optical density values were used in statistical analyses.

### Covariates

Covariate data were extracted from the Multigenerational Register, the Register of Total Population, the Medical birth registry, National Patient Register, the Stockholm Adult psychiatric Outpatient Care Register, the VAL database, and the Integrated Database for Labor Market Research. We considered maternal age at the time of delivery, immigrant status (yes/no), BMI at first antenatal visit, hospitalization for infection in late pregnancy (3^rd^ trimester hospitalizations since these are the most common and more likely to affect markers measured at birth), and psychiatric history before delivery (yes/no); parental educational attainment (highest of mother or father) and income (quintiles of income based on the national distribution, accounting for all sources of income and inflation and adjusting for family size); and children’s sex, birth order, mode of delivery, gestational age birth, size for gestational age, and Apgar score at birth.

### Statistical analyses

Statistical analyses were carried out using STATA/IC^®^ 15.1 (College Station, TX, USA). Distributions of AGA optical densities, both when measured in NDBS and in maternal sera, were right skewed and were log_2_ transformed. We observed operational influences on the distributions of AGA, e.g. yield during protein elution of the NDBS, as well as batch- and plate-to-plate differences for the NDBS (27 plates) as well as for the maternal sera (12 plates). Plate-specific z-scores were therefore calculated from the log_2_ transformed values by subtracting the plate-specific mean from each observation and dividing by the plate-specific standard deviation (SF1).

Cut-offs were then set based on the distribution of these z-scores among controls. No known clinically meaningful cut-off for positive or negative AGA values exists when measured in archival specimens. As it has been suggested that 10-20 % of the normal population carry AGA (20, 21), the cut-off for negative values was set at the 80^th^ percentile of AGA z-scores. Two mutually exclusive categories were created to indicate high values of AGA: from the 80^th^ - >90^th^ percentile value and >90^th^ percentile (z-score).

We employed logistic regression models using this categorical AGA parameterization to determine the odds ratios (OR) and 95% Confidence intervals (CI) for ASD in offspring, comparing each positive category (80->90^th^ percentile and ≥ 90^th^ percentile) to the negative category (<80^th^ percentile). To formally test whether AGA in general was associated with odds of ASD, we used a Wald test of the null-hypothesis that the terms for all AGA categories were equal to zero. Results for this test for association are given as p_association_.

In order to assess the potential for confounding, we examined relationships between covariates and case-control status, and between covariates and AGA levels among controls, using frequency statistics (i.e. the Chi^2^ test). Covariates that were associated to both the exposure and the outcome (p < 0.2) were considered as potential confounders, as were a few additional covariates previously established as strongly associated with the outcome and/or with the exposure (i.e., maternal psychiatric history and mode of delivery).

We compared results from a crude model to the fully adjusted model. Logistic regression models using NDBS categories as the exposure were adjusted for sex, maternal age, maternal BMI, maternal psychiatric history, maternal immigrant status, income, birth order, gestational age at birth and mode of delivery. Logistic regression models using maternal sera categories as the exposure were adjusted for sex, maternal age, maternal immigrant status, maternal psychiatric history, maternal BMI, income, birth order and maternal total IgG. Logistic regression analyses were then repeated with the stratified outcomes based on comorbidities.

We also analyzed AGA as continuous variables to evaluate the relationship between AGA and odds of ASD over the range of AGA measures. We used a restricted cubic spline model with three knots and an AGA Z-score of 0 as the referent to flexibly fit the relationship between AGA and odds of ASD. We used the same scheme for model adjustment as in the categorical models.

In the sibling comparison models, we compared ASD cases to their matched, unaffected siblings using conditional logistic regression models to calculate OR’s and 95% CI for AGA measured in NDBS both in a crude model and adjusted for sex and birth order, factors that are not necessarily shared by siblings.

## Results

### Characteristics of the study samples

Our final sample with analyzed NDBS consisted of 2011 individuals (1090 controls, 921 ASD cases), the sample with paired NDBS-maternal serum specimens consisted of 975 individuals (547 controls, 428 cases), and the sibling sample consisted of 412 individuals (206 unaffected siblings and 206 cases).

ASD cases were more likely to be male, first born, preterm, delivered by CS, to be small or large for gestational age, and to have a lower Apgar score; and were more likely to be born to mothers who were older, who had a higher BMI, who had a psychiatric history, who migrated to Sweden, and who were hospitalized with infection during late pregnancy, as compared to controls (ST2-3).

### Covariates associated with levels of AGA

Maternal age, maternal BMI, sex and gestational age at birth were associated with levels of AGA measured in the NDBS (Table 1) as well as with the outcome (p < 0.2) and hence met the *a priori* criteria for inclusion in the models. This pattern of associations differed for AGA measured in maternal serum where maternal immigration status, family income, and parity were associated with levels of AGA (ST4). Since these co-variates were also associated with the outcome in the larger neonatal sample (p < 0.2), they were included in the models.

**Table 1.** Association of covariates with AGA levels measured in NDBS. Absolute and relative frequencies (n/%) of controls (N = 1,090) in each covariate category are shown according to AGA category (i.e. IgG values < 80^th^, 80-90^th^ or >90^th^ percentile according to the distribution observed among controls) as measured in NDBS.

|  | AGA category |  |  |  |  |  |
| --- | --- | --- | --- | --- | --- | --- |
|  | < 80<sup>th</sup> percentile |  | 80<sup>th</sup> – 90<sup>th</sup> percentile |  | ≥ 90<sup>th</sup> percentile |  |
|  | N | % | N | % | N | % |
| **Sex <sup>a</sup>** |  |  |  |  |  |  |
| Male | 444 | 80.43 | 47 | 8.51 | 61 | 11.05 |
| Female | 428 | 79.55 | 62 | 11.52 | 48 | 8.92 |
| **Maternal age <sup>a</sup>** |  |  |  |  |  |  |
| <25 | 107 | 74.31 | 17 | 11.81 | 20 | 13.89 |
| 25-29 | 234 | 80.14 | 25 | 8.56 | 33 | 11.30 |
| 30-34 | 331 | 84.22 | 35 | 8.91 | 27 | 6.87 |
| 35-39 | 172 | 75.11 | 29 | 12.66 | 28 | 12.23 |
| ≥40 | 28 | 87.50 | 3 | 9.38 | 1 | 1.13 |
| **Maternal BMI <sup>a</sup>** |  |  |  |  |  |  |
| Underweight | 15 | 68.18 | 6 | 27.27 | 1 | 4.55 |
| Normal | 435 | 81.16 | 49 | 9.14 | 52 | 9.70 |
| Overweight | 139 | 75.54 | 28 | 15.22 | 17 | 9.24 |
| Obese | 37 | 75.51 | 5 | 10.20 | 7 | 14.29 |
| Missing | 246 | 82.27 | 21 | 7.02 | 32 | 10.70 |
| **Maternal Psychiatric History** |  |  |  |  |  |  |
| Yes | 291 | 78.86 | 39 | 10.57 | 39 | 10.57 |
| No | 581 | 80.58 | 70 | 9.71 | 70 | 9.71 |
| **Mother born outside Sweden** |  |  |  |  |  |  |
| Yes | 221 | 77.54 | 34 | 11.93 | 30 | 10.53 |
| No | 651 | 80.87 | 75 | 9.32 | 79 | 9.81 |
| **Highest Parental education** |  |  |  |  |  |  |
| <9 years | 128 | 77.58 | 22 | 13.33 | 15 | 9.09 |
| 9-12 years | 386 | 81.61 | 38 | 8.03 | 49 | 10.36 |
| > 12 years | 358 | 79.20 | 49 | 10.84 | 45 | 9.96 |
| **Income Quintile** |  |  |  |  |  |  |
| 1<sup>st</sup> (Lowest) | 118 | 76.62 | 19 | 12.34 | 17 | 11.04 |
| 2<sup>nd</sup> | 192 | 82.05 | 18 | 7.69 | 24 | 10.26 |
| 3<sup>rd</sup> | 185 | 77.73 | 26 | 10.92 | 27 | 11.34 |
| 4<sup>th</sup> | 190 | 82.97 | 22 | 9.61 | 17 | 7.42 |
| 5<sup>th</sup> | 187 | 79.57 | 24 | 10.21 | 24 | 10.21 |
| **Maternal hospitalization with infection (3<sup>rd</sup> trimester)** |  |  |  |  |  |  |
| Yes | 22 | 70.97 | 6 | 19.35 | 3 | 9.68 |
| No | 796 | 79.76 | 99 | 9.92 | 103 | 10.32 |
| Missing | 54 | 88.52 | 4 | 6.56 | 3 | 4.92 |
| **Birth order** |  |  |  |  |  |  |
| 1<sup>st</sup> born | 350 | 79.01 | 46 | 10.38 | 47 | 10.61 |
| 2<sup>nd</sup> born | 307 | 80.16 | 35 | 9.14 | 41 | 10.70 |
| 3<sup>rd</sup> born | 162 | 79.02 | 24 | 11.71 | 19 | 9.27 |
| Missing | 53 | 89.83 | 4 | 6.78 | 2 | 3.39 |
| **Mode of delivery** |  |  |  |  |  |  |
| Unassisted VD | 566 | 79.72 | 71 | 10.00 | 73 | 10.28 |
| Induced, Unassisted VD | 50 | 79.37 | 6 | 9.52 | 7 | 11.11 |
| Assisted VD | 84 | 82.35 | 10 | 9.80 | 8 | 7.84 |
| Elective CS | 47 | 72.31 | 8 | 12.31 | 10 | 15.38 |
| Emergency CS | 72 | 79.12 | 10 | 10.99 | 9 | 9.89 |
| Missing | 53 | 89.83 | 4 | 6.78 | 2 | 3.39 |
| **Gestational age at birth <sup>a</sup>** |  |  |  |  |  |  |
| Term | 703 | 79.52 | 91 | 10.29 | 90 | 10.18 |
| Preterm | 47 | 88.68 | 1 | 1.89 | 5 | 9.43 |
| Post-term | 68 | 73.91 | 13 | 14.13 | 11 | 11.96 |
| Missing | 54 | 88.52 | 4 | 6.56 | 3 | 4.92 |
| **Size for gestational age** |  |  |  |  |  |  |
| Normal | 764 | 79.75 | 98 | 10.23 | 96 | 10.02 |
| Small for GA | 24 | 82.76 | 1 | 3.45 | 4 | 13.79 |
| Large for GA | 27 | 75.00 | 5 | 13.89 | 4 | 11.11 |
| Missing | 57 | 85.07 | 5 | 7.46 | 5 | 7.46 |
| **Apgar score** |  |  |  |  |  |  |
| ≤7 | 9 | 60.00 | 3 | 20.00 | 3 | 20.00 |
| 8 | 26 | 78.79 | 5 | 15.15 | 2 | 6.06 |
| 9 | 81 | 73.64 | 15 | 13.64 | 14 | 12.73 |
| 10 | 691 | 80.44 | 82 | 9.55 | 86 | 10.01 |
| Missing | 65 | 89.04 | 4 | 5.48 | 4 | 5.48 |
| **Total Protein Quartile** |  |  |  |  |  |  |
| 1 | 222 | 25.46 | 30 | 27.52 | 21 | 19.27 |
| 2 | 213 | 24.43 | 30 | 27.52 | 29 | 26.61 |
| 3 | 220 | 25.23 | 25 | 22.94 | 28 | 25.69 |
| 4 | 217 | 24.89 | 24 | 22.02 | 21 | 28.44 |

Associations between the listed covariates and AGA exposure were calculated using the Chi2 test, missing values not included.
<sup>a</sup> The covariate is associated with the exposure at $p < 0.2$
Abbreviations: ADHD = Attention deficit hyperactivity disorder, AGA = Antigliadin antibodies, ASD = Autism spectrum disorders, BMI = Body Mass Index, CS = Cesarean section, ID = Intellectual disability, GA = Gestational age, NDBS = Neonatal dried blood spots, VD = Vaginal delivery

### Antigliadin antibody levels and ASD in offspring

In maternal sera and NDBS respectively, 16.35 % and 18.78 % of cases had high levels of AGA compared to 20 % of the respective controls, as defined by the cut-off (Table 2).

**Table 2.** The absolute and relative frequencies of study individuals according to AGA categories, for AGA measured in maternal sera and in NDBS.

|  | <80<sup>th</sup> |  | 80-90<sup>th</sup> |  | >90<sup>th</sup> |  | Total |  | *p*<sup>a</sup> |
| --- | --- | --- | --- | --- | --- | --- | --- | --- | --- |
|  | N | % | N | % | N | % | N | % |  |
| AGA measured in maternal sera |  |  |  |  |  |  |  |  |  |
| Controls | 437 | 79.89 | 55 | 10.05 | 55 | 10.05 | 547 | 100 |  |
| Cases | 358 | 83.64 | 33 | 7.71 | 37 | 8.64 | 428 | 100 | 0.304 |
| ASD only | 106 | 80.30 | 13 | 9.85 | 13 | 9.85 | 132 | 100 | 0.994 |
| ASD with ID | 129 | 89.58 | 8 | 5.56 | 7 | 4.86 | 144 | 100 | 0.026 |
| ASD with ADHD | 123 | 80.92 | 12 | 7.89 | 17 | 11.18 | 152 | 100 | 0.690 |
| AGA measured in NDBS |  |  |  |  |  |  |  |  |  |
| Controls | 872 | 80.00 | 109 | 10.00 | 109 | 10.00 | 1090 | 100 |  |
| Cases | 748 | 81.22 | 99 | 10.75 | 74 | 8.03 | 921 | 100 | 0.289 |
| ASD only | 223 | 79.36 | 31 | 11.03 | 27 | 9.61 | 281 | 100 | 0.870 |
| ASD with ID | 262 | 82.39 | 37 | 11.64 | 19 | 5.97 | 318 | 100 | 0.074 |
| ASD with ADHD | 263 | 81.68 | 31 | 9.63 | 28 | 8.70 | 322 | 100 | 0.757 |
| AGA measured in NDBS (Sibling comparison) |  |  |  |  |  |  |  |  |  |
| Controls | 155 | 75.24 | 20 | 9.71 | 31 | 15.05 | 206 | 100 |  |
| Cases | 171 | 83.00 | 17 | 8.25 | 18 | 8.74 | 206 | 100 | 0.107 |
| ASD only | 51 | 79.69 | 6 | 9.38 | 7 | 10.94 | 64 | 100 | 0.698 |
| ASD with ID | 77 | 84.62 | 9 | 9.89 | 5 | 5.49 | 91 | 100 | 0.065 |
| ASD with ADHD | 43 | 84.31 | 2 | 3.92 | 6 | 11.76 | 51 | 100 | 0.310 |

<sup>a</sup> P-value for the Chi<sup>2</sup> test, used to test the association between AGA category and case-control status.
Abbreviations: AGA = Antigliadin antibodies, ASD = Autism spectrum disorders, ADHD = Attention deficit hyperactive disorder, ID = Intellectual disability

In the sibling sample, 16.99 % of cases, and 24.76 % of the unaffected siblings were positive for AGA (Table 2). AGA levels in the 80^th^-90^th^ percentile category in maternal sera were not associated with any of the outcomes (Table 3, SF2). Levels ≥ 90^th^ percentile, were associated with significantly decreased odds of ASD with comorbid ID (OR 0.43, 95% CI 0.19–0.97), though the association was attenuated in the fully adjusted model (OR 0.55, 95% CI 0.24-1.29). No significant associations were seen for the other outcomes studied.

**Table 3.**
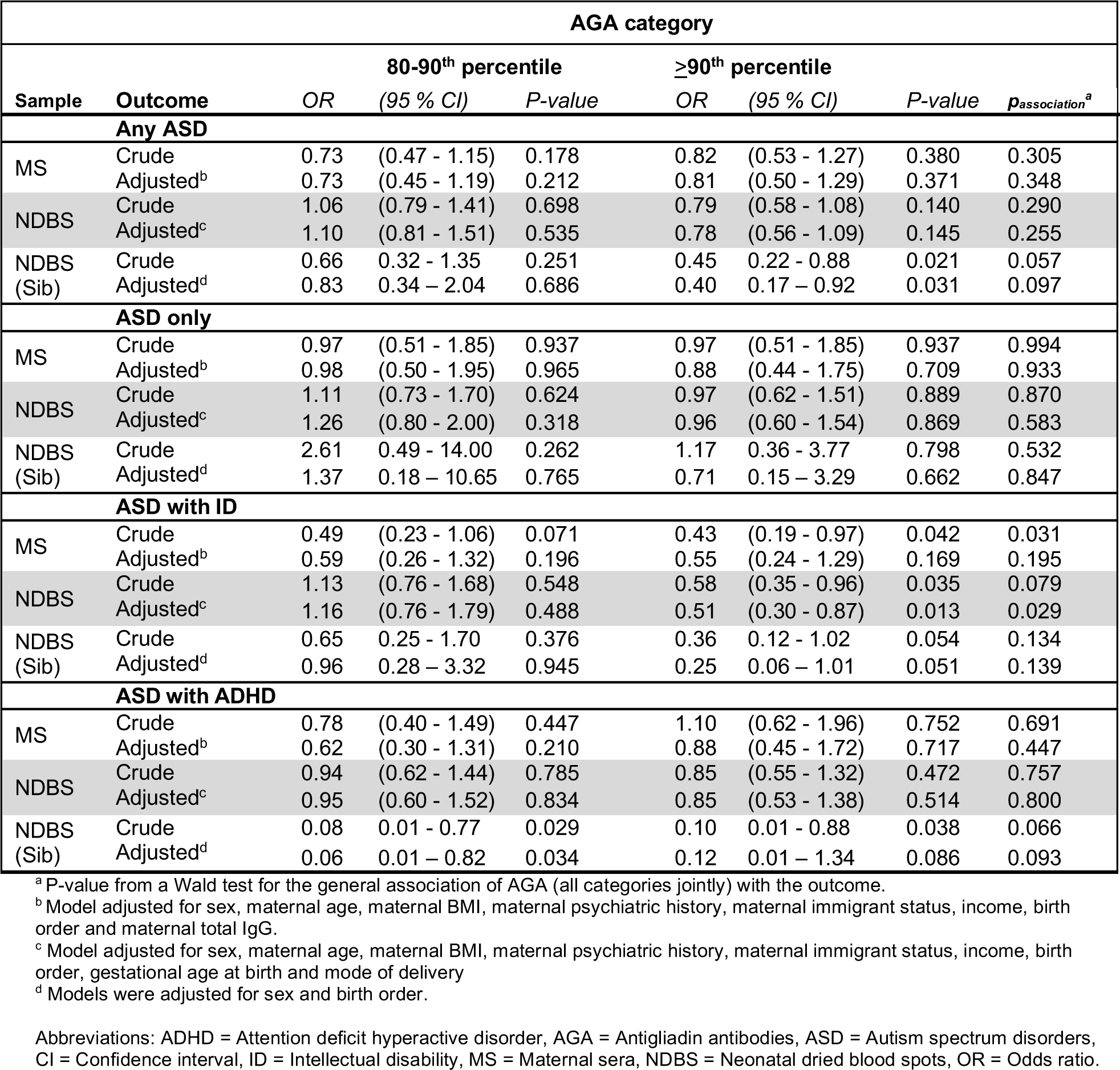
Maternal AGA levels during pregnancy and offspring odds of ASD. Crude and fully adjusted OR’s and the 95 % CI associated with higher categories of maternal AGA levels, when measured in MS and NDBS respectively, using values below the 80^th^ percentile as a referent. Results are shown for any ASD diagnosis and for diagnoses stratified by the presence of co-occurring ID and ADHD.

AGA levels in the 80^th^-90^th^ percentile category measured in NDBS were not associated with any of the outcomes (Table 3, SF2). Levels ≥90^th^ percentile were associated with significantly decreased odds of ASD with comorbid ID in a crude model (OR 0.58, 95 % CI 0.35–0.96), as well as a fully adjusted model (OR 0.51, 95 % CI 0.30–0.87). No significant associations were seen for the other outcomes, though point estimates were consistently below the null value, with the exception for ASD without comorbid ID or ADHD (i.e. ASD “only”).

### AGA and odds of ASD in the sibling comparison

High levels of AGA (≥ 90^th^ percentile), measured in NDBS, were significantly associated with decreased odds of ASD as a combined diagnostic group and with ASD with ADHD (both 80^th^-90^th^ percentile and ≥90^th^ percentile) levels in sibling comparison analyses (Table 3, SF3).

Adjusting for sex and birth order did not affect the association with ASD as a combined diagnostic group, but attenuated the association between AGA and ASD with ADHD. The estimated OR for ASD with comorbid ID was substantially below one, albeit not statistically significant.

### Considering AGA as a continuous variable

We observed a pattern of a lower odds of ASD (as a combined diagnostic group) and of ASD with ID with increasing AGA levels, both when measured in maternal sera and in NDBS (Figure 2).

**Figure 2.**
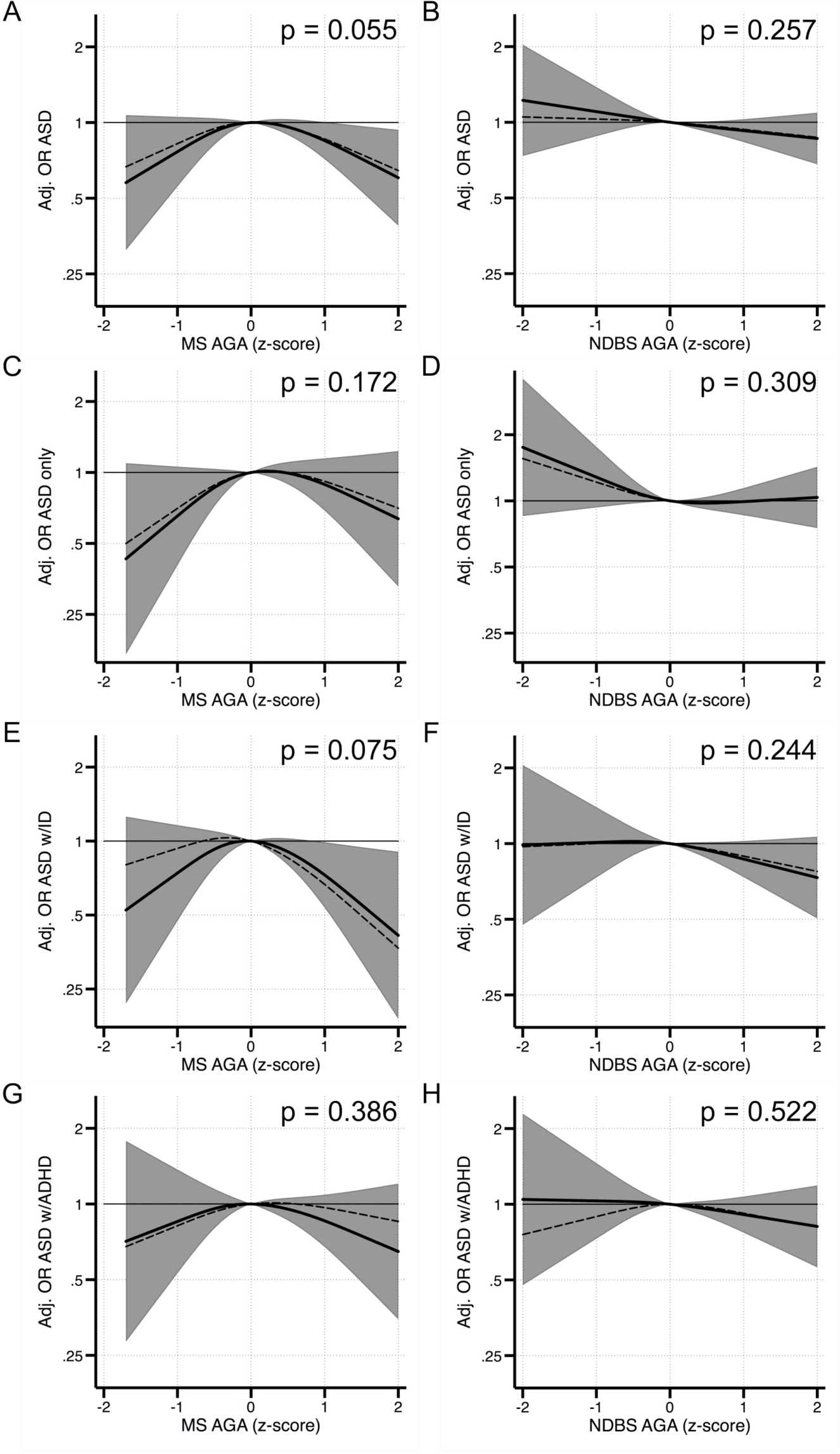
Maternal AGA and odds of ASD in the offspring. Each panel displays the odds of each outcome according to AGA z-score, flexibly fit using restricted cubic spline models with three knots and a z-score=0 as the referent. The dashed line represents the unadjusted estimate of the relationship between AGA and odds of each outcome. The solid line represents the fully adjusted model. Models for AGA in MS were adjusted for sex, maternal age, maternal BMI, maternal psychiatric history, maternal immigrant status, income, maternal total IgG and birth order. Models for AGA in NDBS were adjusted for sex, maternal age, maternal BMI, maternal psychiatric history, maternal immigrant status, income, birth order, gestational age at birth and mode of delivery. The gray bands represent the 95% confidence interval for the fully adjusted model. P-values are shown for a Wald test with a null hypothesis that all AGA spline terms were jointly equal to zero, as a test of whether AGA was generally associated with the outcome. Results are shown for odds associated with AGA in MS (on the left) and AGA in NDBS (on the right) for the outcomes of any ASD diagnosis (**A- B**), ASD without co-occurring ID or ADHD (“ASD only”; **C-D**), ASD with co-occurring ID (**E-F**), and ASD with co-occurring ADHD (**G-H**). Abbreviations; ADHD = Attention deficit hyperactive disorder, ASD = Autism spectrum disorder, AGA = Antigliadin antibodies, BMI = Body mass index, CI = Confidence interval, ID = Intellectual disability, MS = Maternal sera, NDBS = Neonatal dried blood spots, OR = Odds ratio

## Discussion

We found that high levels (≥90^th^ percentile) of maternal AGA during pregnancy, measured in maternal sera and in NDBS, were associated with decreased odds of ASD with comorbid ID in the offspring. For ASD with comorbid ADHD, point estimates tended to be below one, indicative of a decrease in offspring odds for ASD. These observed associations were mirrored in the sibling study. Taken together, our findings suggest that *high* levels of maternal AGA during pregnancy, may be protective against ASD, particularly ASD with comorbid ID, in the offspring.

This is the first study to date to investigate the association between maternal AGA during pregnancy and the later development of ASD in offspring. Considering the numerous studies reporting high levels of AGA in individuals already diagnosed with ASD (12–14) and in individuals diagnosed with other adult-onset neuropsychiatric disorders (22, 23), our results are unexpected.

On the other hand, our results are consistent with results from a few studies investigating the relationship between maternal IgG antibodies directed against other specific antigens, such as those derived from *Toxoplasma gondii* (*T. gondii*), and ASD in offspring (24–26) in which high levels of maternal IgG were associated with decreased offspring risk of ASD.

### Potential mechanism and implications for ASD pathogenesis

Our results support the notion that the immune system is of potential etiological relevance in ASD and provides novel data indicating that maternally-derived IgG antibodies directed towards gluten components might be implicated.

The mechanisms underlying this association are uncertain. Maternal IgG antibodies are actively transferred across the human placenta at an increasing rate towards the end of pregnancy to provide the neonate with passive immunity against antigens in the maternal environment (27). Hypothetically, the observed association between high levels IgG antibodies and lowered risk of ASD in neonatal blood might be explained by impaired transplacental transfer of maternal antibodies in ASD pregnancies (25). However, our study does not support this hypothesis since similar associations was observed also in maternal sera obtained earlier in that same in pregnancy.

It has also been hypothesized that risk associated with lower levels of maternal IgG represent an overall dysfunctional maternal immunity resulting in reduced levels of circulating IgG among mothers to children with ASD (24, 25). This possibility is an unlikely explanation for our findings, since adjustment for maternal total IgG levels, measured in maternal sera, did not attenuate the associations between AGA and ASD observed maternal sera. Moreover, we found no association between maternal total IgG quintiles and case-control status. Thus, we find no evidence that the observed associations for the maternal IgG antigliadin antibodies would simply be a proxy for total maternal IgG levels.

Though we were able to adjust for a wide range of potential confounders, including maternal total IgG, and attempted to account for shared familial factors in a sibling study design, residual confounding cannot be excluded. Considering that these antibodies presumably reflect maternal dietary exposure to gluten, they may indicate a role for the nutritional status of the mother. Maternal nutrition is a factor associated with ASD that might plausibly correlate with the presence of AGA. For example, maternal iron deficiency during the early stages of pregnancy has been associated with an increased risk of ASD in offspring (28). Cereals, including wheat, constitute an important source of iron in Sweden (29) and the reduced ASD risk associated with AGA may reflect wheat consumption and adequate iron supplies in a subgroup of women. We have no data on dietary habits among mothers in the current study sample. Adjusting for maternal BMI (admittedly a poor proxy for dietary intake/nutritional status) did not markedly attenuate the associations.

Also, adjustment for maternal migration status attenuated the associations. It is plausible that dietary habits and the likelihood of developing AGA varies between different countries and ethnicities. The prevalence of ASD, particularly ASD with co-morbid ID, varies by maternal country of origin (30), which may introduce confounding not fully accounted for in the present study. However, we observed similar patterns of associations in the sibling comparison models that would be expected to account for this particular source of bias.

Finally, we consider that AGA may more directly play a protective role in the etiology of ASD with comorbid ID. First, it is possible that these maternally derived antibodies provide the newborn with protection against harmful effects of gluten during the perinatal period. Gliadin has been detected in human breast milk (31) and may be introduced with complementary foods from four months of age (32). It could thus be speculated that exposure to gliadin in early life, without protective immunity transferred from the mother, might be harmful. An effect of gluten on CNS function in humans has been described (33), based on the fact that digestion of gluten results in the generation of stable peptides, some of which have been reported to have opioid activity (33). Additional studies need to be performed to explore this possibility.

Second, it is possible that transferred AGA cross-react with and thereby clears some other harmful exogenous antigen(s) and in that way protects the newborn against ASD. This intriguing possibility should also be explored in order to identify any such antigens.

Thirdly, antibodies directed towards gliadin have been reported to exhibit cross-reactivity with cerebellar proteins (34, 35). It may thus be possible that circulating maternal AGA, by binding to gliadin, protects the newborn from generating high levels of such auto-reactive antibodies themselves. We cannot further explore such mechanisms in the current dataset.

### Strengths and limitation

We used a validated ASD case ascertainment procedure set in the context of a healthcare system where developmental screenings are routinely carried out free of charge. The comparison group was randomly selected from population-based birth cohorts, increasing the generalizability. All data were prospectively obtained without regard to later diagnoses. A major advantage is that we were able to analyze both neonatal and paired maternal specimens. Though IgG in neonatal circulation are presumed to consist of maternally derived IgG, the maternal sera analysis provides a technically less challenging sample for the assessment of maternal humoral immunity. These paired samples also allowed us to account for the role played by the placental transfer. On the same note, we were able to analyze and adjust for levels of maternal total IgG, which minimizes the possibility that our results reflect maternal humoral immunity in general. Finally, we used different statistical approaches, analyzing AGA as both a categorical and a continuous variable, as well as using a sibling comparison design, all with similar results, strengthening the interpretation of our findings.

With regard to limitations, the number of study individuals exceeding the cut-offs in the categorical analyses was small, especially for the stratified analyses and the sibling analyses, limiting the power of those analyses. Moreover, as previously discussed, our results may still be influenced by residual confounding. However, the similar pattern of associations observed in the sibling analysis suggest that shared familial environmental or genetic factors are unlikely to explain observed associations. The largest limitation to our study is the limited knowledge regarding how to interpret the levels AGA measured in general, including the lack of clinically meaningful cut-offs when analyzing these antibodies.

## Conclusion

Taken together, our findings consistently suggest that high levels of maternal AGA during pregnancy, are protective against ASD in the offspring, particularly against ASD with comorbid ID. As this is the first study to examine maternal AGA and ASD risk, further research is necessary to test the replicability of our results and to elucidate how higher levels of AGA during the perinatal period may be linked to lower risk of ASD.

## Acknowledgements

This work was supported by grants from the Swedish Research Council (grant numbers 2016-01477, 2012-2264 and 523-2010-1052 [to CD]), and the Stanley Medical Research Institute [to HK and RY]. The funders had no role in the design and conduct of the study; collection, management, analysis, and interpretation of the data; preparation, review, or approval of the manuscript; or decision to submit the manuscript for publication. We thank Shuojia Yang for her technical expertise in performing the laboratory immunoassays.

## Supplementary materials

### Supplemental Figures

**SF1.**
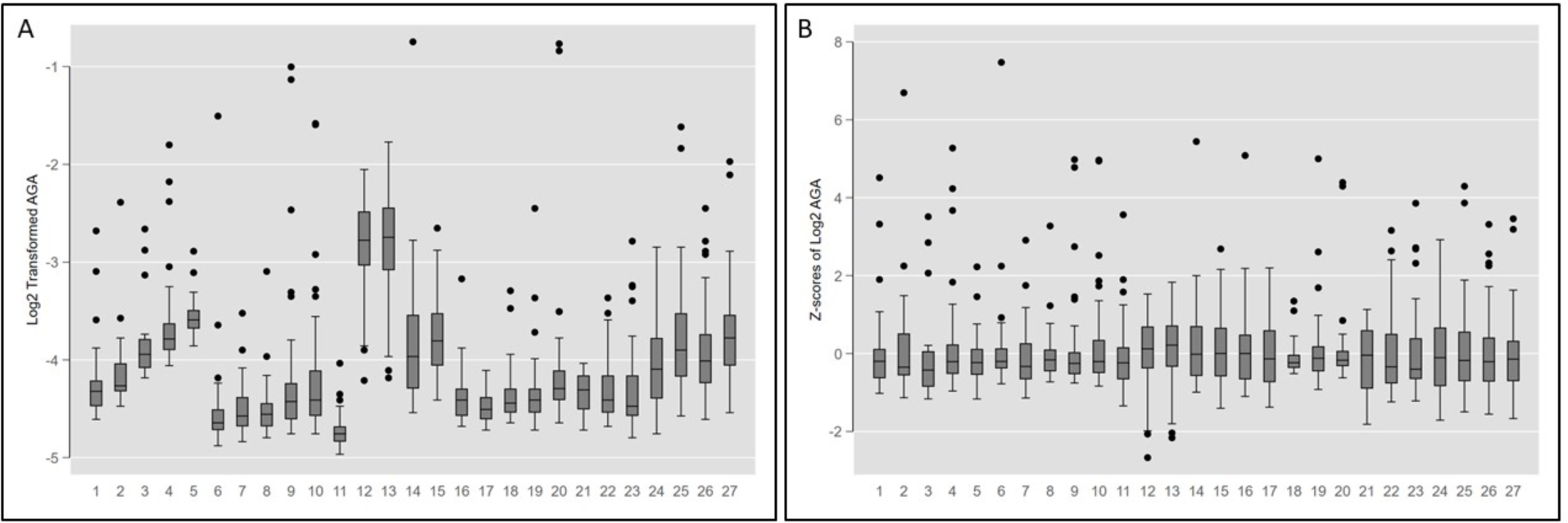
Log_2_ transformed AGA and Z-scores of log_2_ transformed AGA analyzed in NDBS, by analysis plate

**SF2.**
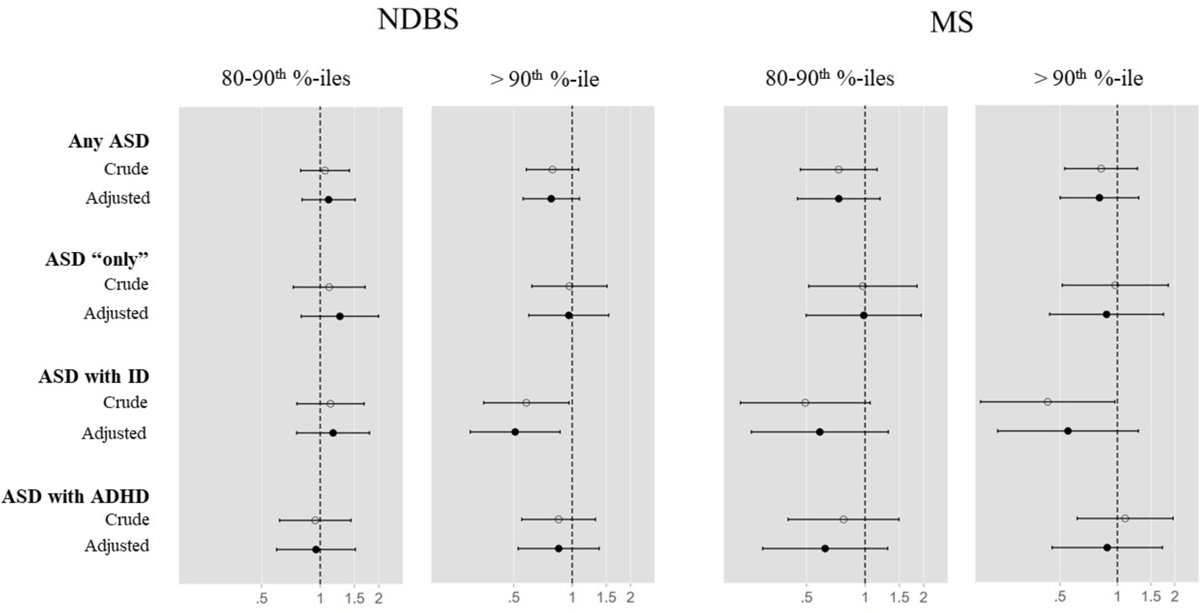
Maternal AGA levels during pregnancy and offspring odds of ASD.

**SF3.**
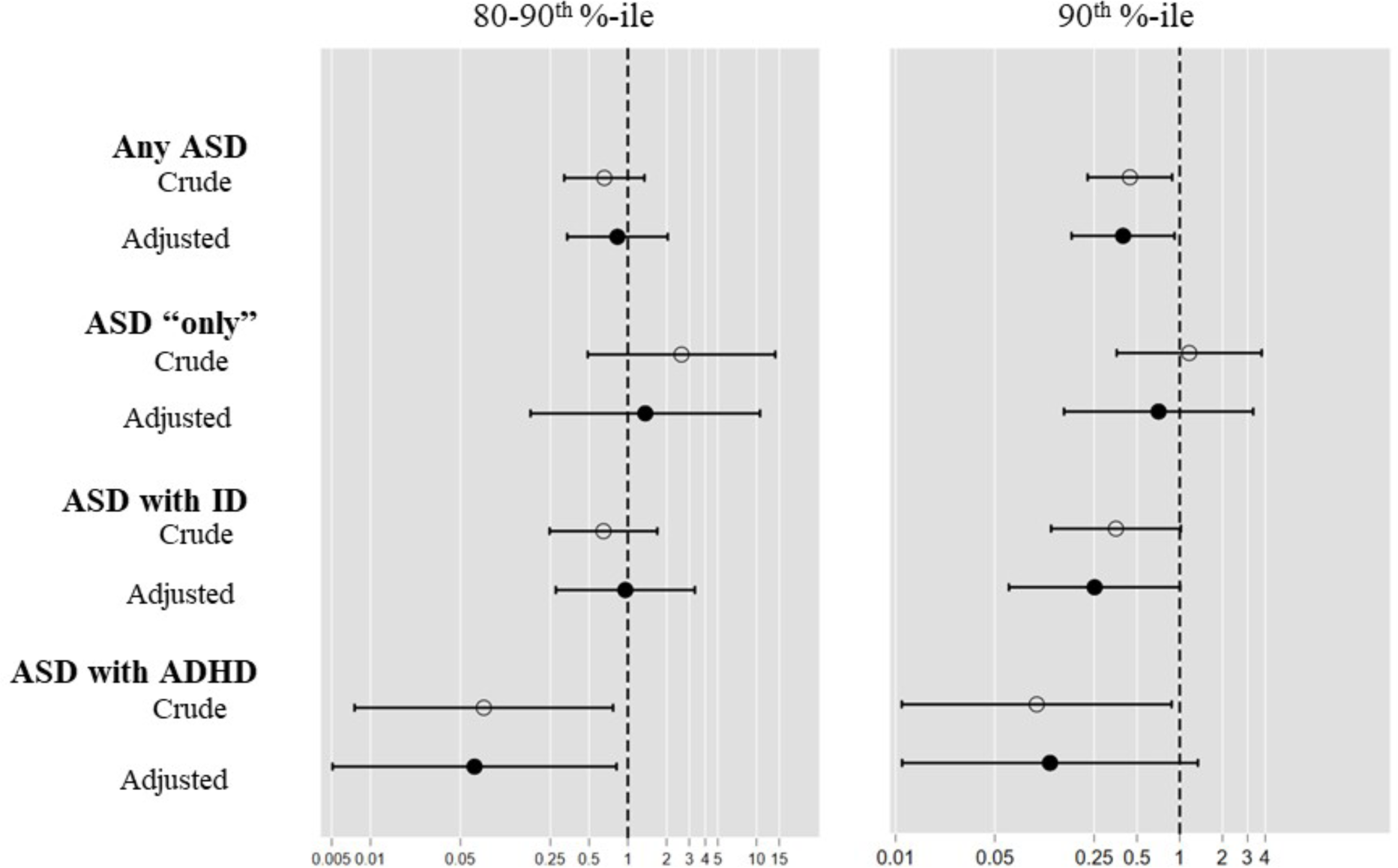
Maternal AGA levels during pregnancy and offspring odds of ASD in the sibling sample.

### Supplemental Tables

**ST1.** Diagnostic codes used in the ascertainment of ASD, ADHD, and ID in the Stockholm Youth Cohort.

| **Coding system** | **Autism Spectrum Disorders (ASD)** | **Attention Deficit/Hyperactivity Disorder (ADHD)** | **Intellectual Disability (ID)** |
| --- | --- | --- | --- |
| ICD-10<sup>a,b,c</sup> | F84 | F90 | F70-F79 |
| DSM-IV<sup>b</sup> | 299 | 314 | 317-319 |
| Other | Habilitation Register<sup>d</sup> | Prescription Drug Register<sup>e</sup>:<br>methylphenidate [N06BA04] or atomoxetine [N06BA09] | Habilitation Register<sup>d</sup> |

<sup>a</sup> The National Patient Register (NPR): including inpatient care from 1973, outpatient physician visits in specialist care from 1997, and children and adolescent psychiatry from 2006.
<sup>b</sup> Stockholm Clinical Database for Child and Adolescent Psychiatry: DSM-IV was used until 2008; ICD-10 was used thereafter.
<sup>c</sup> VAL register: Stockholm county health care databases including in- and outpatient care, regardless of specialty/primary health care.
<sup>d</sup> The Habilitation Register (HAB): Provides data on utilization of Stockholm County Habilitation services according to type of disability, such as pervasive developmental disorders and intellectual disability (recorded as present or absent). HAB register was used until 2008; since 2008 it has been included in the VAL databases as ICD-coded diagnoses (see C above).
<sup>e</sup> The Prescription Drug Register (PDR) contains data on medications dispensed to the entire population in Sweden since 1 July 2005. Receipt of a prescription for ADHD medications is a useful proxy for an ADHD diagnosis, as Swedish medical guidelines mandate that ADHD medications should only be prescribed by a psychiatric specialist and after other (non-pharmacological) interventions have failed.

**ST2.** Characteristics of the NDBS analytical sample, including unaffected siblings.

|  | Controls<br>N = 1,090 |  | ASD cases<br>N = 921 |  | ASD with ID<br>N = 318 |  | ASD with<br>ADHD<br>N = 322 |  | ASD "only"<br>N = 281 |  | Unaffected<br>Siblings<br>N = 206 |  |
| --- | --- | --- | --- | --- | --- | --- | --- | --- | --- | --- | --- | --- |
|  | N | % | N | % | N | % | N | % | N | % | N | % |
| **Sex <sup>b</sup>** |  |  |  |  |  |  |  |  |  |  |  |  |
| Males | 552 | 50.64 | 707 | 76.76 | 234 | 73.58 | 252 | 78.26 | 221 | 78.65 | 89 | 43.20 |
| Females | 538 | 49.36 | 214 | 23.24 | 84 | 26.42 | 70 | 21.74 | 60 | 21.35 | 117 | 56.80 |
| **Maternal Age <sup>b</sup>** |  |  |  |  |  |  |  |  |  |  |  |  |
| <25 | 144 | 13.21 | 115 | 12.49 | 50 | 15.72 | 40 | 12.42 | 25 | 8.90 | 35 | 16.99 |
| 25-29 | 292 | 26.79 | 305 | 33.12 | 106 | 33.33 | 118 | 36.65 | 81 | 28.83 | 68 | 33.01 |
| 30-34 | 393 | 36.06 | 291 | 31.60 | 86 | 27.04 | 102 | 31.68 | 103 | 36.65 | 77 | 37.38 |
| 35-39 | 229 | 21.01 | 173 | 18.78 | 56 | 17.61 | 56 | 17.39 | 61 | 21.71 | 20 | 9.71 |
| ≥40 | 32 | 2.94 | 37 | 4.02 | 20 | 6.29 | 6 | 1.86 | 11 | 3.91 | 6 | 2.91 |
| **Maternal BMI <sup>b</sup>** |  |  |  |  |  |  |  |  |  |  |  |  |
| Underweight | 22 | 2.02 | 22 | 2.39 | 8 | 2.52 | 10 | 3.11 | 4 | 1.42 | 4 | 1.94 |
| Normal | 536 | 49.17 | 362 | 39.31 | 128 | 40.25 | 110 | 34.16 | 124 | 44.13 | 95 | 46.12 |
| Overweight | 184 | 16.88 | 165 | 17.92 | 61 | 19.18 | 62 | 19.25 | 42 | 14.95 | 41 | 19.90 |
| Obese | 49 | 4.50 | 77 | 8.36 | 25 | 7.86 | 35 | 10.87 | 17 | 6.05 | 15 | 7.28 |
| Missing | 299 | 27.43 | 295 | 32.02 | 96 | 30.19 | 105 | 32.61 | 94 | 33.45 | 51 | 24.76 |
| **Maternal Psychiatric History <sup>b</sup>** |  |  |  |  |  |  |  |  |  |  |  |  |
|  | 369 | 33.85 | 439 | 47.67 | 129 | 40.57 | 177 | 54.97 | 133 | 47.33 | 89 | 43.20 |
| **Mother born outside Sweden <sup>a</sup>** |  |  |  |  |  |  |  |  |  |  |  |  |
|  | 285 | 26.15 | 276 | 29.97 | 152 | 47.80 | 60 | 18.63 | 64 | 22.78 | 73 | 35.44 |
| **Highest Parental education <sup>c, a</sup>** |  |  |  |  |  |  |  |  |  |  |  |  |
| <9 years | 165 | 15.14 | 166 | 18.02 | 73 | 22.96 | 59 | 18.32 | 34 | 12.10 | 41 | 19.90 |
| 9-12 years | 473 | 43.39 | 411 | 44.63 | 140 | 44.03 | 155 | 48.14 | 116 | 41.28 | 92 | 44.66 |
| > 12 years | 452 | 41.47 | 342 | 37.13 | 104 | 32.70 | 108 | 33.54 | 130 | 46.26 | 72 | 34.95 |
| **Income Quintile <sup>c, b</sup>** |  |  |  |  |  |  |  |  |  |  |  |  |
| 1<sup>st</sup> (Lowest) | 154 | 14.13 | 122 | 13.25 | 64 | 20.13 | 29 | 9.01 | 29 | 10.32 | 29 | 14.08 |
| 2<sup>nd</sup> | 234 | 21.47 | 239 | 25.95 | 91 | 28.62 | 82 | 25.47 | 66 | 23.49 | 48 | 23.30 |
| 3<sup>rd</sup> | 238 | 21.83 | 214 | 23.24 | 68 | 21.38 | 84 | 26.09 | 62 | 22.06 | 45 | 21.84 |
| 4<sup>th</sup> | 229 | 21.01 | 205 | 22.26 | 64 | 20.13 | 82 | 25.47 | 59 | 21.00 | 44 | 21.36 |
| 5<sup>th</sup> | 235 | 21.56 | 139 | 15.09 | 30 | 9.43 | 45 | 13.98 | 64 | 22.78 | 39 | 18.93 |
| **Maternal hospitalization with infection (3<sup>rd</sup> trimester) <sup>b</sup>** |  |  |  |  |  |  |  |  |  |  |  |  |
| Yes | 31 | 2.84 | 43 | 4. 67 | 19 | 5.97 | 11 | 3.42 | 13 | 4.63 | 7 | 3.40 |
| Missing | 61 | 5.60 | 70 | 7. 60 | 23 | 7.23 | 28 | 8.70 | 19 | 6.76 | 6 | 2.91 |
| **Birth order <sup>b</sup>** |  |  |  |  |  |  |  |  |  |  |  |  |
| 1st born | 443 | 40.64 | 416 | 45.17 | 121 | 38.05 | 158 | 49.07 | 137 | 48.75 | 101 | 49.03 |
| 2nd born | 383 | 35.14 | 289 | 31.38 | 114 | 35.85 | 87 | 27.02 | 88 | 31.32 | 68 | 33.01 |
| 3<sup>rd</sup> born | 205 | 18.81 | 148 | 16.07 | 60 | 18.87 | 49 | 15.22 | 39 | 13.88 | 31 | 15.05 |
| Missing | 59 | 5.41 | 68 | 7.38 | 23 | 7.23 | 28 | 8.70 | 17 | 6.05 | 6 | 2.91 |
| **Mode of Delivery <sup>b</sup>** |  |  |  |  |  |  |  |  |  |  |  |  |
| Unassisted VD | 710 | 65.14 | 530 | 57.55 | 173 | 54.40 | 180 | 55.90 | 177 | 62.99 | 131 | 63.59 |
| Induced, Unassisted VD | 63 | 5.78 | 71 | 7.71 | 23 | 7.23 | 30 | 9.32 | 18 | 6.41 | 10 | 4. 85 |
| Assisted VD | 102 | 9.36 | 89 | 9.66 | 36 | 11.32 | 29 | 9.01 | 24 | 8.54 | 20 | 9.71 |
| Elective CS | 65 | 5.96 | 71 | 7.71 | 26 | 8.18 | 21 | 6.52 | 24 | 8.54 | 15 | 7.28 |
| Emergency CS | 91 | 8.35 | 92 | 9.99 | 37 | 11.64 | 34 | 10.56 | 21 | 7.47 | 24 | 11.65 |
| Missing | 59 | 5.41 | 68 | 7.38 | 23 | 7.23 | 28 | 8.70 | 17 | 6.05 | 6 | 2.91 |
| **Gestational Age <sup>a</sup>** |  |  |  |  |  |  |  |  |  |  |  |  |
| Term | 884 | 81.10 | 720 | 78.18 | 246 | 77.36 | 250 | 77.64 | 224 | 79.72 | 175 | 84.95 |
| Preterm | 53 | 4.86 | 63 | 6.84 | 23 | 7.23 | 21 | 6.52 | 19 | 6. 76 | 8 | 3.88 |
| Post-term | 92 | 8.44 | 69 | 7.49 | 26 | 8.18 | 23 | 7.14 | 20 | 7.12 | 17 | 8.25 |
| Missing | 61 | 5.60 | 69 | 7.49 | 23 | 7.23 | 28 | 8.70 | 18 | 6.41 | 6 | 2.91 |
| **Size for Gestational Age <sup>a</sup>** |  |  |  |  |  |  |  |  |  |  |  |  |
| Normal | 958 | 87.89 | 769 | 83.50 | 271 | 85.22 | 258 | 80.12 | 240 | 85.41 | 185 | 89.81 |
| Small for GA | 29 | 2.66 | 29 | 3.15 | 14 | 4.40 | 10 | 3.11 | 5 | 1.78 | 9 | 4.37 |
| Large for GA | 36 | 3.30 | 43 | 4.67 | 8 | 2.52 | 20 | 6.21 | 15 | 5.34 | 6 | 2.91 |
| Missing | 67 | 6.15 | 80 | 8.69 | 25 | 7.86 | 34 | 10.56 | 21 | 7.47 | 6 | 2.91 |
| **Apgar score (5 minutes) <sup>a</sup>** |  |  |  |  |  |  |  |  |  |  |  |  |
| ≤7 | 15 | 1. 38 | 25 | 2. 71 | 19 | 5.97 | 3 | 0.93 | 3 | 1.07 | 5 | 2.43 |
| 8 | 33 | 3.03 | 36 | 3.91 | 20 | 6.29 | 10 | 3.11 | 6 | 2.14 | 9 | 4.37 |
| 9 | 110 | 10.09 | 101 | 10.97 | 35 | 11.01 | 36 | 11.18 | 30 | 10.68 | 25 | 12.14 |
| 10 | 859 | 78.81 | 682 | 74.05 | 216 | 67.92 | 243 | 75.47 | 223 | 79.36 | 160 | 77.67 |
| Missing | 73 | 6.70 | 77 | 8.36 | 28 | 8.81 | 30 | 9.32 | 19 | 6.76 | 7 | 3.40 |

| Total Protein Quartile |  |  |  |  |  |  |  |  |  |  |  |  |
| --- | --- | --- | --- | --- | --- | --- | --- | --- | --- | --- | --- | --- |
| 1 | 273 | 25.05 | 235 | 25.52 | 91 | 28.62 | 78 | 24.22 | 66 | 23.49 | 50 | 24.27 |
| 2 | 272 | 24.95 | 201 | 21.82 | 64 | 20.13 | 66 | 20.50 | 71 | 25.27 | 37 | 17.96 |
| 3 | 273 | 25.05 | 230 | 24.97 | 80 | 25.16 | 78 | 24.22 | 72 | 25.62 | 64 | 21.07 |
| 4 | 272 | 24.95 | 255 | 27.69 | 83 | 26.10 | 100 | 31.06 | 72 | 25.62 | 55 | 26.70 |

Associations between the listed covariates and case-control status were calculated using the Chi<sup>2</sup> test, missing values not included.
<sup>a</sup> The covariate is associated with the outcome at $p < 0.2$
<sup>b</sup> The covariate is associated with the outcome at $p < 0.05$
<sup>c</sup> Missing data for 2 ASD cases and 1 unaffected sibling.
Abbreviations: ADHD = Attention deficit hyperactivity disorder, ASD = Autism spectrum disorders, BMI = Body Mass Index, CS = Cesarean section, ID = Intellectual disability, GA = Gestational age, NDBS = Neonatal dried blood spots, VD = Vaginal delivery.

**ST3.**
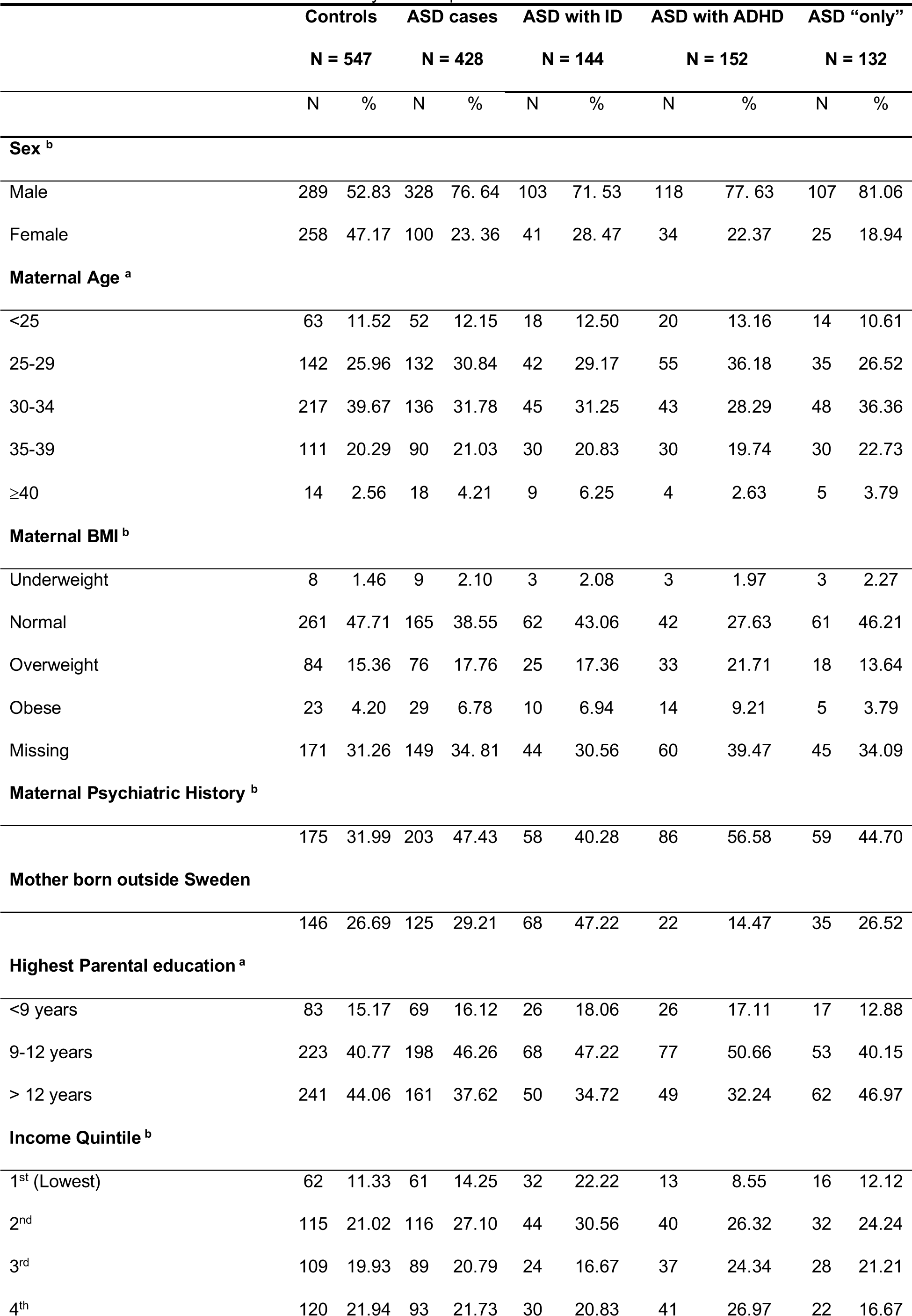
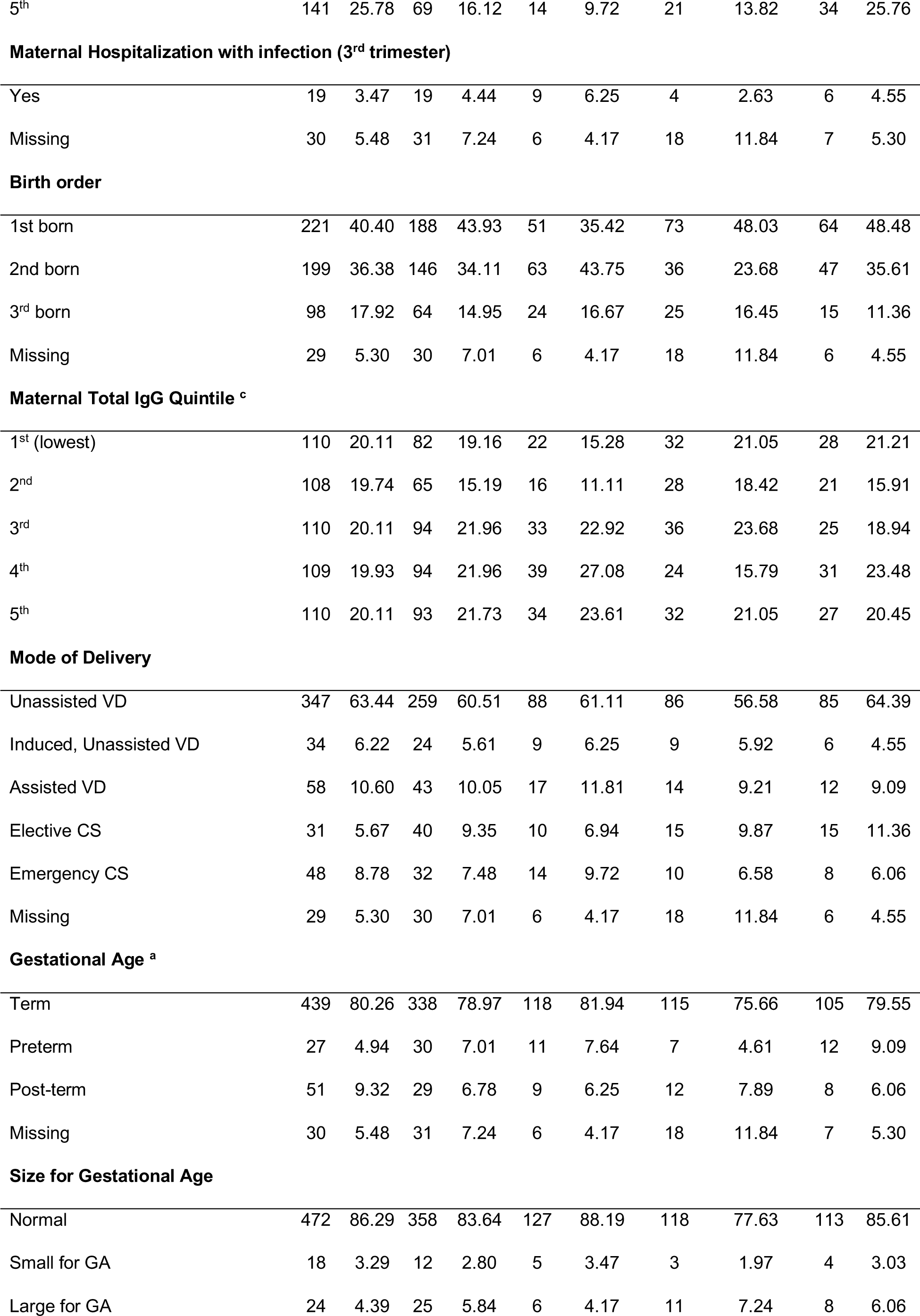
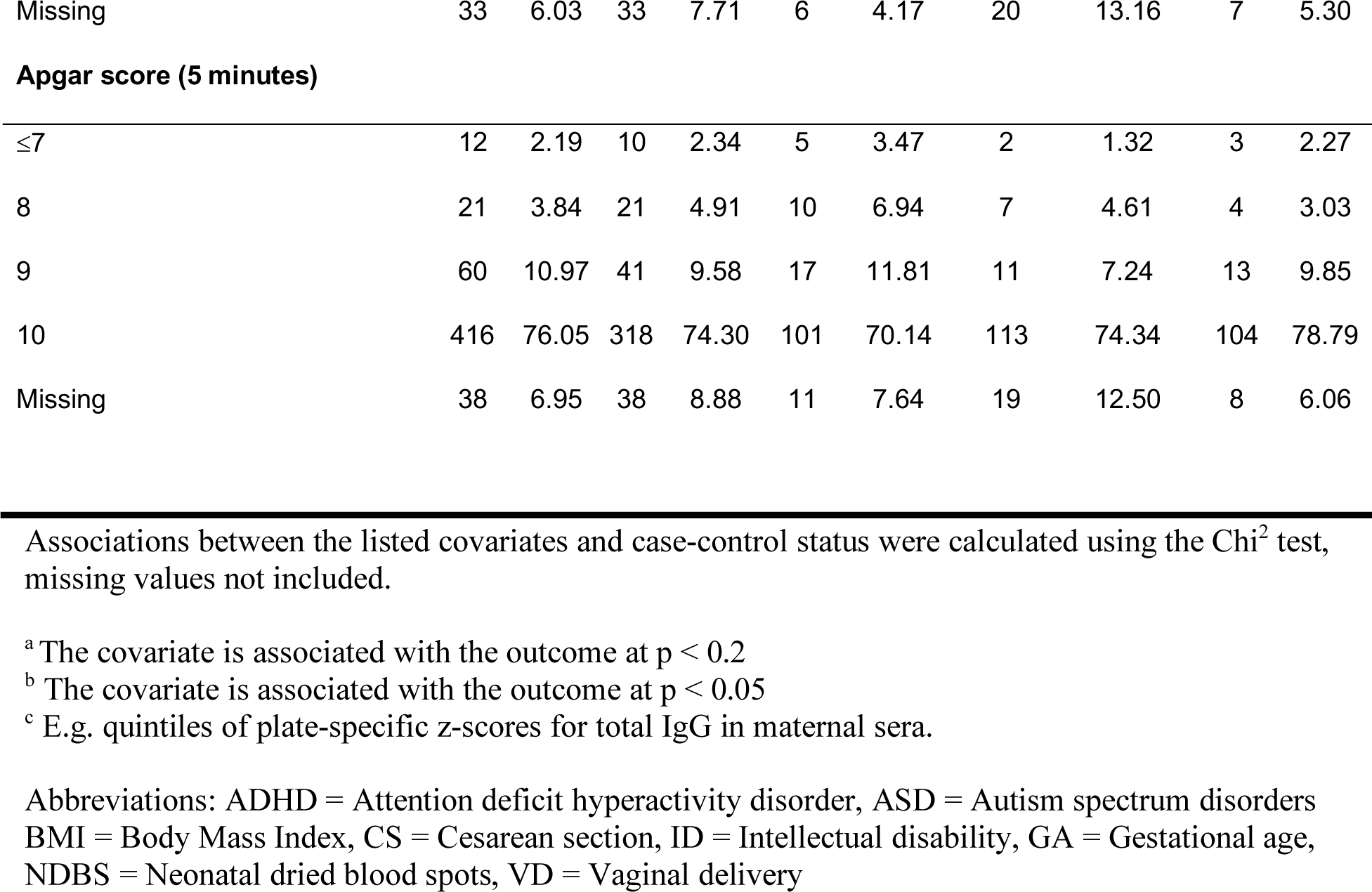
Characteristics of the maternal sera analytical sample.

**ST3. Characteristics of the maternal sera analytical sample.**
|  | **Controls** |  | **ASD cases** |  | **ASD with ID** |  | **ASD with ADHD** |  | **ASD “only”** |  |
| --- | --- | --- | --- | --- | --- | --- | --- | --- | --- | --- |
|  | **N = 547** |  | **N = 428** |  | **N = 144** |  | **N = 152** |  | **N = 132** |  |
|  | N | % | N | % | N | % | N | % | N | % |
| **Sex <sup>b</sup>** |  |  |  |  |  |  |  |  |  |  |
| Male | 289 | 52.83 | 328 | 76.64 | 103 | 71.53 | 118 | 77.63 | 107 | 81.06 |
| Female | 258 | 47.17 | 100 | 23.36 | 41 | 28.47 | 34 | 22.37 | 25 | 18.94 |
| **Maternal Age <sup>a</sup>** |  |  |  |  |  |  |  |  |  |  |
| <25 | 63 | 11.52 | 52 | 12.15 | 18 | 12.50 | 20 | 13.16 | 14 | 10.61 |
| 25-29 | 142 | 25.96 | 132 | 30.84 | 42 | 29.17 | 55 | 36.18 | 35 | 26.52 |
| 30-34 | 217 | 39.67 | 136 | 31.78 | 45 | 31.25 | 43 | 28.29 | 48 | 36.36 |
| 35-39 | 111 | 20.29 | 90 | 21.03 | 30 | 20.83 | 30 | 19.74 | 30 | 22.73 |
| ≥40 | 14 | 2.56 | 18 | 4.21 | 9 | 6.25 | 4 | 2.63 | 5 | 3.79 |
| **Maternal BMI <sup>b</sup>** |  |  |  |  |  |  |  |  |  |  |
| Underweight | 8 | 1.46 | 9 | 2.10 | 3 | 2.08 | 3 | 1.97 | 3 | 2.27 |
| Normal | 261 | 47.71 | 165 | 38.55 | 62 | 43.06 | 42 | 27.63 | 61 | 46.21 |
| Overweight | 84 | 15.36 | 76 | 17.76 | 25 | 17.36 | 33 | 21.71 | 18 | 13.64 |
| Obese | 23 | 4.20 | 29 | 6.78 | 10 | 6.94 | 14 | 9.21 | 5 | 3.79 |
| Missing | 171 | 31.26 | 149 | 34.81 | 44 | 30.56 | 60 | 39.47 | 45 | 34.09 |
| **Maternal Psychiatric History <sup>b</sup>** |  |  |  |  |  |  |  |  |  |  |
|  | 175 | 31.99 | 203 | 47.43 | 58 | 40.28 | 86 | 56.58 | 59 | 44.70 |
| **Mother born outside Sweden** |  |  |  |  |  |  |  |  |  |  |
|  | 146 | 26.69 | 125 | 29.21 | 68 | 47.22 | 22 | 14.47 | 35 | 26.52 |
| **Highest Parental education <sup>a</sup>** |  |  |  |  |  |  |  |  |  |  |
| <9 years | 83 | 15.17 | 69 | 16.12 | 26 | 18.06 | 26 | 17.11 | 17 | 12.88 |
| 9-12 years | 223 | 40.77 | 198 | 46.26 | 68 | 47.22 | 77 | 50.66 | 53 | 40.15 |
| > 12 years | 241 | 44.06 | 161 | 37.62 | 50 | 34.72 | 49 | 32.24 | 62 | 46.97 |
| **Income Quintile <sup>b</sup>** |  |  |  |  |  |  |  |  |  |  |
| 1<sup>st</sup> (Lowest) | 62 | 11.33 | 61 | 14.25 | 32 | 22.22 | 13 | 8.55 | 16 | 12.12 |
| 2<sup>nd</sup> | 115 | 21.02 | 116 | 27.10 | 44 | 30.56 | 40 | 26.32 | 32 | 24.24 |
| 3<sup>rd</sup> | 109 | 19.93 | 89 | 20.79 | 24 | 16.67 | 37 | 24.34 | 28 | 21.21 |
| 4<sup>th</sup> | 120 | 21.94 | 93 | 21.73 | 30 | 20.83 | 41 | 26.97 | 22 | 16.67 |
| 5<sup>th</sup> | 141 | 25.78 | 69 | 16.12 | 14 | 9.72 | 21 | 13.82 | 34 | 25.76 |
| **Maternal Hospitalization with infection (3<sup>rd</sup> trimester)** |  |  |  |  |  |  |  |  |  |  |
| Yes | 19 | 3.47 | 19 | 4.44 | 9 | 6.25 | 4 | 2.63 | 6 | 4.55 |
| Missing | 30 | 5.48 | 31 | 7.24 | 6 | 4.17 | 18 | 11.84 | 7 | 5.30 |
| **Birth order** |  |  |  |  |  |  |  |  |  |  |
| 1st born | 221 | 40.40 | 188 | 43.93 | 51 | 35.42 | 73 | 48.03 | 64 | 48.48 |
| 2nd born | 199 | 36.38 | 146 | 34.11 | 63 | 43.75 | 36 | 23.68 | 47 | 35.61 |
| 3<sup>rd</sup> born | 98 | 17.92 | 64 | 14.95 | 24 | 16.67 | 25 | 16.45 | 15 | 11.36 |
| Missing | 29 | 5.30 | 30 | 7.01 | 6 | 4.17 | 18 | 11.84 | 6 | 4.55 |
| **Maternal Total IgG Quintile <sup>c</sup>** |  |  |  |  |  |  |  |  |  |  |
| 1<sup>st</sup> (lowest) | 110 | 20.11 | 82 | 19.16 | 22 | 15.28 | 32 | 21.05 | 28 | 21.21 |
| 2<sup>nd</sup> | 108 | 19.74 | 65 | 15.19 | 16 | 11.11 | 28 | 18.42 | 21 | 15.91 |
| 3<sup>rd</sup> | 110 | 20.11 | 94 | 21.96 | 33 | 22.92 | 36 | 23.68 | 25 | 18.94 |
| 4<sup>th</sup> | 109 | 19.93 | 94 | 21.96 | 39 | 27.08 | 24 | 15.79 | 31 | 23.48 |
| 5<sup>th</sup> | 110 | 20.11 | 93 | 21.73 | 34 | 23.61 | 32 | 21.05 | 27 | 20.45 |
| **Mode of Delivery** |  |  |  |  |  |  |  |  |  |  |
| Unassisted VD | 347 | 63.44 | 259 | 60.51 | 88 | 61.11 | 86 | 56.58 | 85 | 64.39 |
| Induced, Unassisted VD | 34 | 6.22 | 24 | 5.61 | 9 | 6.25 | 9 | 5.92 | 6 | 4.55 |
| Assisted VD | 58 | 10.60 | 43 | 10.05 | 17 | 11.81 | 14 | 9.21 | 12 | 9.09 |
| Elective CS | 31 | 5.67 | 40 | 9.35 | 10 | 6.94 | 15 | 9.87 | 15 | 11.36 |
| Emergency CS | 48 | 8.78 | 32 | 7.48 | 14 | 9.72 | 10 | 6.58 | 8 | 6.06 |
| Missing | 29 | 5.30 | 30 | 7.01 | 6 | 4.17 | 18 | 11.84 | 6 | 4.55 |
| **Gestational Age <sup>a</sup>** |  |  |  |  |  |  |  |  |  |  |
| Term | 439 | 80.26 | 338 | 78.97 | 118 | 81.94 | 115 | 75.66 | 105 | 79.55 |
| Preterm | 27 | 4.94 | 30 | 7.01 | 11 | 7.64 | 7 | 4.61 | 12 | 9.09 |
| Post-term | 51 | 9.32 | 29 | 6.78 | 9 | 6.25 | 12 | 7.89 | 8 | 6.06 |
| Missing | 30 | 5.48 | 31 | 7.24 | 6 | 4.17 | 18 | 11.84 | 7 | 5.30 |
| **Size for Gestational Age** |  |  |  |  |  |  |  |  |  |  |
| Normal | 472 | 86.29 | 358 | 83.64 | 127 | 88.19 | 118 | 77.63 | 113 | 85.61 |
| Small for GA | 18 | 3.29 | 12 | 2.80 | 5 | 3.47 | 3 | 1.97 | 4 | 3.03 |
| Large for GA | 24 | 4.39 | 25 | 5.84 | 6 | 4.17 | 11 | 7.24 | 8 | 6.06 |
| Missing | 33 | 6.03 | 33 | 7.71 | 6 | 4.17 | 20 | 13.16 | 7 | 5.30 |
| **Apgar score (5 minutes)** |  |  |  |  |  |  |  |  |  |  |
| ≤7 | 12 | 2.19 | 10 | 2.34 | 5 | 3.47 | 2 | 1.32 | 3 | 2.27 |
| 8 | 21 | 3.84 | 21 | 4.91 | 10 | 6.94 | 7 | 4.61 | 4 | 3.03 |
| 9 | 60 | 10.97 | 41 | 9.58 | 17 | 11.81 | 11 | 7.24 | 13 | 9.85 |
| 10 | 416 | 76.05 | 318 | 74.30 | 101 | 70.14 | 113 | 74.34 | 104 | 78.79 |
| Missing | 38 | 6.95 | 38 | 8.88 | 11 | 7.64 | 19 | 12.50 | 8 | 6.06 |

Associations between the listed covariates and case-control status were calculated using the Chi<sup>2</sup> test, missing values not included.
<sup>a</sup> The covariate is associated with the outcome at $p < 0.2$
<sup>b</sup> The covariate is associated with the outcome at $p < 0.05$
<sup>c</sup> E.g. quintiles of plate-specific z-scores for total IgG in maternal sera.
Abbreviations: ADHD = Attention deficit hyperactivity disorder, ASD = Autism spectrum disorders BMI = Body Mass Index, CS = Cesarean section, ID = Intellectual disability, GA = Gestational age, NDBS = Neonatal dried blood spots, VD = Vaginal delivery

**ST4.**
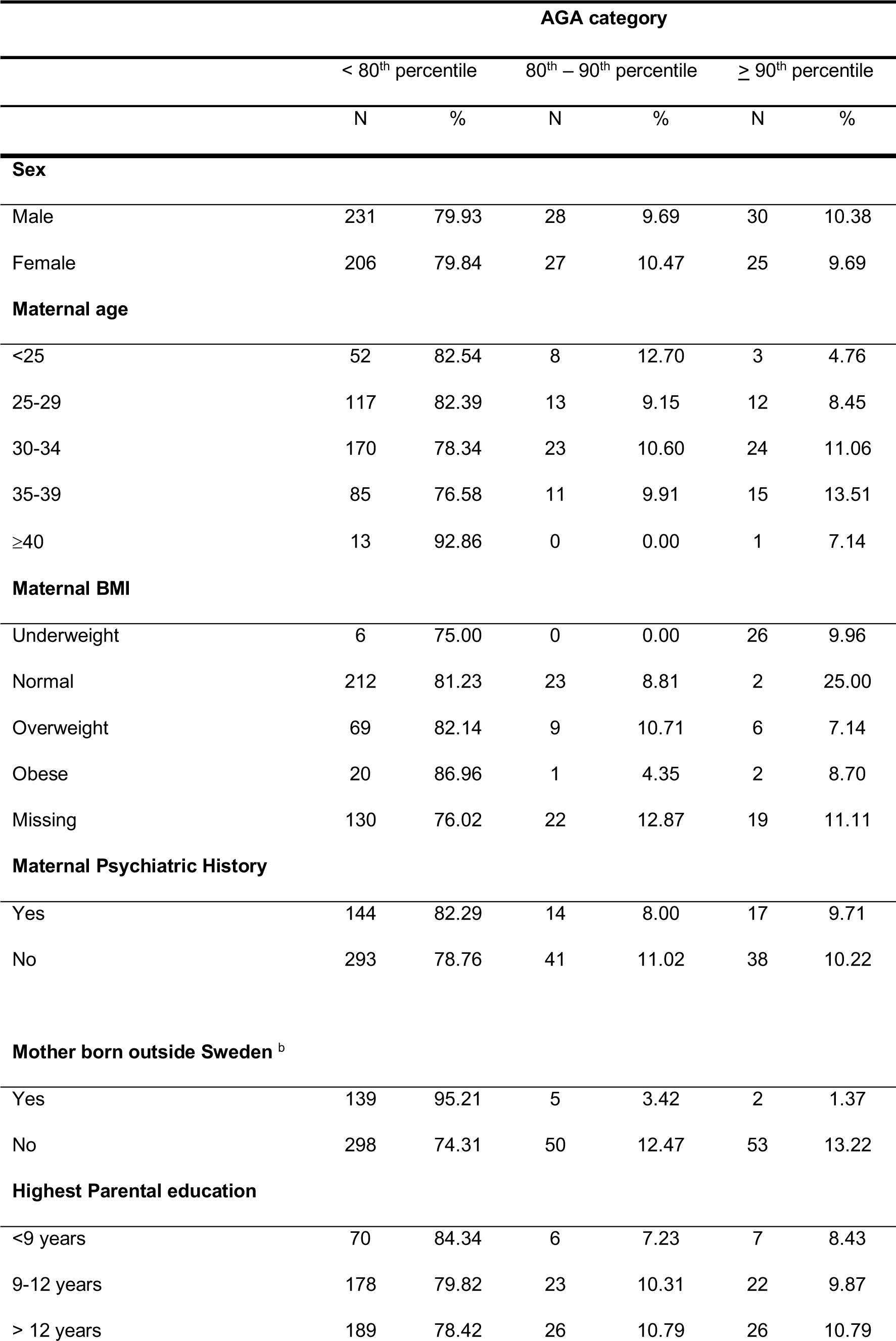
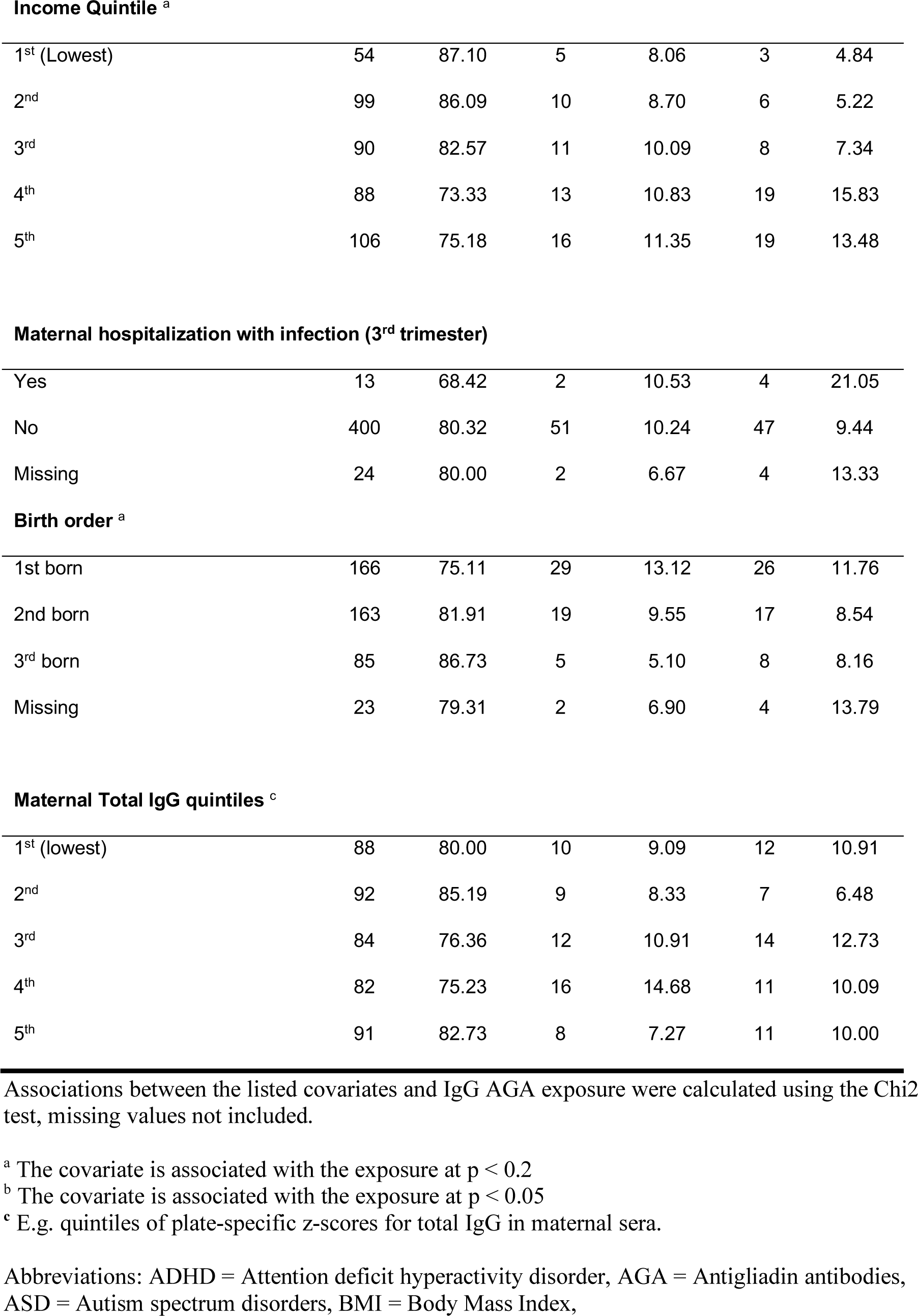
Association of covariates with AGA levels measured in maternal sera.

|  | AGA category |  |  |  |  |  |
| --- | --- | --- | --- | --- | --- | --- |
|  | < 80<sup>th</sup> percentile |  | 80<sup>th</sup> – 90<sup>th</sup> percentile |  | ≥ 90<sup>th</sup> percentile |  |
|  | N | % | N | % | N | % |
| **Sex** |  |  |  |  |  |  |
| Male | 231 | 79.93 | 28 | 9.69 | 30 | 10.38 |
| Female | 206 | 79.84 | 27 | 10.47 | 25 | 9.69 |
| **Maternal age** |  |  |  |  |  |  |
| <25 | 52 | 82.54 | 8 | 12.70 | 3 | 4.76 |
| 25-29 | 117 | 82.39 | 13 | 9.15 | 12 | 8.45 |
| 30-34 | 170 | 78.34 | 23 | 10.60 | 24 | 11.06 |
| 35-39 | 85 | 76.58 | 11 | 9.91 | 15 | 13.51 |
| ≥40 | 13 | 92.86 | 0 | 0.00 | 1 | 7.14 |
| **Maternal BMI** |  |  |  |  |  |  |
| Underweight | 6 | 75.00 | 0 | 0.00 | 26 | 9.96 |
| Normal | 212 | 81.23 | 23 | 8.81 | 2 | 25.00 |
| Overweight | 69 | 82.14 | 9 | 10.71 | 6 | 7.14 |
| Obese | 20 | 86.96 | 1 | 4.35 | 2 | 8.70 |
| Missing | 130 | 76.02 | 22 | 12.87 | 19 | 11.11 |
| **Maternal Psychiatric History** |  |  |  |  |  |  |
| Yes | 144 | 82.29 | 14 | 8.00 | 17 | 9.71 |
| No | 293 | 78.76 | 41 | 11.02 | 38 | 10.22 |
| **Mother born outside Sweden <sup>b</sup>** |  |  |  |  |  |  |
| Yes | 139 | 95.21 | 5 | 3.42 | 2 | 1.37 |
| No | 298 | 74.31 | 50 | 12.47 | 53 | 13.22 |
| **Highest Parental education** |  |  |  |  |  |  |
| <9 years | 70 | 84.34 | 6 | 7.23 | 7 | 8.43 |
| 9-12 years | 178 | 79.82 | 23 | 10.31 | 22 | 9.87 |
| > 12 years | 189 | 78.42 | 26 | 10.79 | 26 | 10.79 |
| **Income Quintile <sup>a</sup>** |  |  |  |  |  |  |
| 1<sup>st</sup> (Lowest) | 54 | 87.10 | 5 | 8.06 | 3 | 4.84 |
| 2<sup>nd</sup> | 99 | 86.09 | 10 | 8.70 | 6 | 5.22 |
| 3<sup>rd</sup> | 90 | 82.57 | 11 | 10.09 | 8 | 7.34 |
| 4<sup>th</sup> | 88 | 73.33 | 13 | 10.83 | 19 | 15.83 |
| 5<sup>th</sup> | 106 | 75.18 | 16 | 11.35 | 19 | 13.48 |
| **Maternal hospitalization with infection (3<sup>rd</sup> trimester)** |  |  |  |  |  |  |
| Yes | 13 | 68.42 | 2 | 10.53 | 4 | 21.05 |
| No | 400 | 80.32 | 51 | 10.24 | 47 | 9.44 |
| Missing | 24 | 80.00 | 2 | 6.67 | 4 | 13.33 |
| **Birth order <sup>a</sup>** |  |  |  |  |  |  |
| 1st born | 166 | 75.11 | 29 | 13.12 | 26 | 11.76 |
| 2nd born | 163 | 81.91 | 19 | 9.55 | 17 | 8.54 |
| 3<sup>rd</sup> born | 85 | 86.73 | 5 | 5.10 | 8 | 8.16 |
| Missing | 23 | 79.31 | 2 | 6.90 | 4 | 13.79 |
| **Maternal Total IgG quintiles <sup>c</sup>** |  |  |  |  |  |  |
| 1<sup>st</sup> (lowest) | 88 | 80.00 | 10 | 9.09 | 12 | 10.91 |
| 2<sup>nd</sup> | 92 | 85.19 | 9 | 8.33 | 7 | 6.48 |
| 3<sup>rd</sup> | 84 | 76.36 | 12 | 10.91 | 14 | 12.73 |
| 4<sup>th</sup> | 82 | 75.23 | 16 | 14.68 | 11 | 10.09 |
| 5<sup>th</sup> | 91 | 82.73 | 8 | 7.27 | 11 | 10.00 |

Associations between the listed covariates and IgG AGA exposure were calculated using the Chi2 test, missing values not included.
<sup>a</sup> The covariate is associated with the exposure at $p < 0.2$
<sup>b</sup> The covariate is associated with the exposure at $p < 0.05$
<sup>c</sup> E.g. quintiles of plate-specific z-scores for total IgG in maternal sera.
Abbreviations: ADHD = Attention deficit hyperactivity disorder, AGA = Antigliadin antibodies, ASD = Autism spectrum disorders, BMI = Body Mass Index,

